# Punishment history biases corticothalamic responses to motivationally-significant stimuli

**DOI:** 10.1101/2020.04.06.027888

**Authors:** Federica Lucantonio, Zhixiao Su, Anna J. Chang, Bilal A. Bari, Jeremiah Y. Cohen

## Abstract

Making predictions about future rewards or punishments is fundamental to adaptive behavior. These processes are influenced by prior experience. For example, prior exposure to aversive stimuli or stressors changes behavioral responses to negative- and positive-value predictive cues. Here, we demonstrate a role for medial prefrontal cortex (mPFC) neurons projecting to the paraventricular nucleus of the thalamus (PVT; mPFC→PVT) in this process. We found that a history of punishments negatively biased behavioral responses to motivationally-relevant stimuli in mice and that this negative bias was associated with hyperactivity in mPFC→PVT neurons during exposure to those cues. Furthermore, artificially mimicking this hyperactive response with selective optogenetic excitation of the same pathway recapitulated the punishment-induced negative behavioral bias. Together, our results highlight how information flow within the mPFC→PVT circuit is critical for making predictions about imminent motivationally-relevant outcomes as a function of prior experience.

## Introduction

Effective decision making requires anticipating biologically significant outcomes associated with environmental stimuli. It also requires balancing the goals of a decision—for instance, acquiring a reward or avoiding a punishment—with certainty about the outcome of the decision. For example, when the outcome of a decision to approach or avoid a stimulus is ambiguous, the nervous system must weigh the cost of receiving a punishment, or missing out on a reward, with the benefit of obtaining the reward, or avoiding the punishment.

Many decisions are influenced by background emotional state. For example, both positive and negative mood affect decision making (Deldin and Levin, 1986; Wright and Bower, 1992; Bechara et al., 2000; Hockey et al., 2000; Dolan, 2002; Harding et al., 2004). This background state can be driven by prior experience. In particular, prior exposure to aversive stimuli or stressors changes behavioral responses to ambiguous stimuli (Harding et al., 2004; Boleij et al., 2012; Rygula et al., 2014). How the balance between competing behaviors is weighed in the brain or how prior experience with an environment shifts this balance is still poorly understood.

The medial prefrontal cortex (mPFC) is critical for regulating cue-mediated behaviors in both appetitive and aversive domains (Ishikawa et al., 2008; Burgos-Robles et al., 2009; Sotres-Bayon and Quirk, 2010; Amemori and Graybiel, 2012; Burgos-Robles et al., 2013; Courtin et al., 2014; Sangha et al., 2014; Sparta et al., 2014; Burgos-Robles et al., 2017; Otis et al., 2017). Behavioral manifestations of appetitive and aversive conditioning correlate with changes in neural activity within the mPFC (Burgos-Robles et al., 2009; Peters et al., 2009; Amemori and Graybiel, 2012; Burgos-Robles et al., 2013; Moorman and Aston-Jones, 2015) and pharmacological or optogenetic manipulations of mPFC alter both reward-seeking and fear-related behaviors (Morgan and LeDoux, 1995; Blum et al., 2006; Corcoran and Quirk, 2007; Sierra-Mercado et al., 2011; Sangha et al., 2014; Sparta et al., 2014; Bari et al., 2019).

The mPFC has dense projections to subcortical structures involved in motivated behavior, including the paraventricular thalamus (PVT; Vertes, 2004; Li and Kirouac, 2012). Like the mPFC, PVT is recruited by cues or contexts previously associated with rewarding or aversive outcomes (Beck and Fibiger, 1995; Schiltz et al., 2007; Yasoshima et al., 2007; Choi et al., 2010; Igelstrom et al., 2010; Haight and Flagel, 2014; Hsu et al., 2014; Do-Monte et al., 2015; Kirouac, 2015; Matzeu et al., 2015; Penzo et al., 2015; Li et al., 2016; Zhu et al., 2016; Do-Monte et al., 2017). PVT neurons are activated by multiple forms of stressors (Chastrette et al., 1991; Sharp et al., 1991; Cullinan et al., 1995; Bubser and Deutch, 1999; Spencer et al., 2004) and coordinate behavioral responses to stress (Hsu et al., 2014; Do-Monte et al., 2015; Penzo et al., 2015; Zhu et al., 2016; Do-Monte et al., 2017; Beas et al., 2018). On the other hand, under conditions of opposing emotional valence, PVT plays a role in multiple forms of stimulus-reward learning and PVT neurons have been reported to show reward-modulated responses (Schiltz et al., 2005; Igelstrom et al., 2010; Martin-Fardon and Boutrel, 2012; James and Dayas, 2013; Browning et al., 2014; Haight and Flagel, 2014; Li et al., 2016; Choi et al., 2019). Activity in mPFC neurons projecting to the PVT also suppresses both the acquisition and expression of conditioned reward seeking (Otis et al., 2017).

Taken together, these studies place the mPFC to PVT projection in a unique position to integrate information about positive and negative motivationally-relevant cues and translate it into adaptive behavioral responses. How these projection-specific prefrontal neurons regulate behavioral responses in ambiguous settings and how their neural activity may be altered upon presentation of a punishment is unknown.

To address these questions, we trained mice on a go/no-go discrimination task in which sweet- and bitter-predicting odor cues together with mixtures of varying proportions of those cues were concurrently presented to the mice and recorded extracellularly in single-cell mPFC and in identified mPFC→PVT neurons. We then tested whether optogenetically stimulating mPFC→PVT projections recapitulated the observed neuron-behavior relationships in those settings.

## Results

### Punishment history negatively biases behavioral responses to motivationally-significant stimuli

To assess the effect of punishment history on decisions about motivationally-significant outcomes, we developed a go/no-go discrimination task in head-fixed mice consisting of four phases: conditioning, probe test, reversal, and a second probe test (Figure 1A). In the conditioning phase, four odor cues (A, B, C and D, counterbalanced) were presented. Odor A predicted an appetitive sweet solution (3 *µ*l of 5% sucrose water). Odor B predicted an aversive bitter solution (3 *µ*l of 10 mM denatonium water). Odor C was associated with no reinforcement. Odor D predicted a punishment (an unavoidable air puff delivered to the mouse’s right eye) in mice assigned to the air puff group and was associated with no reinforcement in mice assigned to the no air puff group. Each behavioral trial began with an odor (1 s; conditioned stimulus, CS), followed by a 1-s delay and an outcome (unconditioned stimulus, US). Mice showed essentially binary responses to cues that predicted sucrose or denatonium. That is, they licked in anticipation of sucrose and did not in anticipation of denatonium. Thus, we designed a probe test, in which, in addition to the four conditioning odors, we measured behavioral responses to ambiguous stimuli. We designed parametrically-varying mixtures of stimuli between appetitive and aversive solutions. We exposed mice to unreinforced mixtures of varying proportions of odors A and B: 85%A/15%B, 50%A/50%B, 15%A/85%B. To ensure that behavioral responses to ambiguous stimuli were not driven by a mouse’s preference for a particular odor, after completion of the probe test, cue-outcome associations for odors A and B were reversed, and each mouse was re-tested in a second set of probe stimuli.

**Fig. 1.**
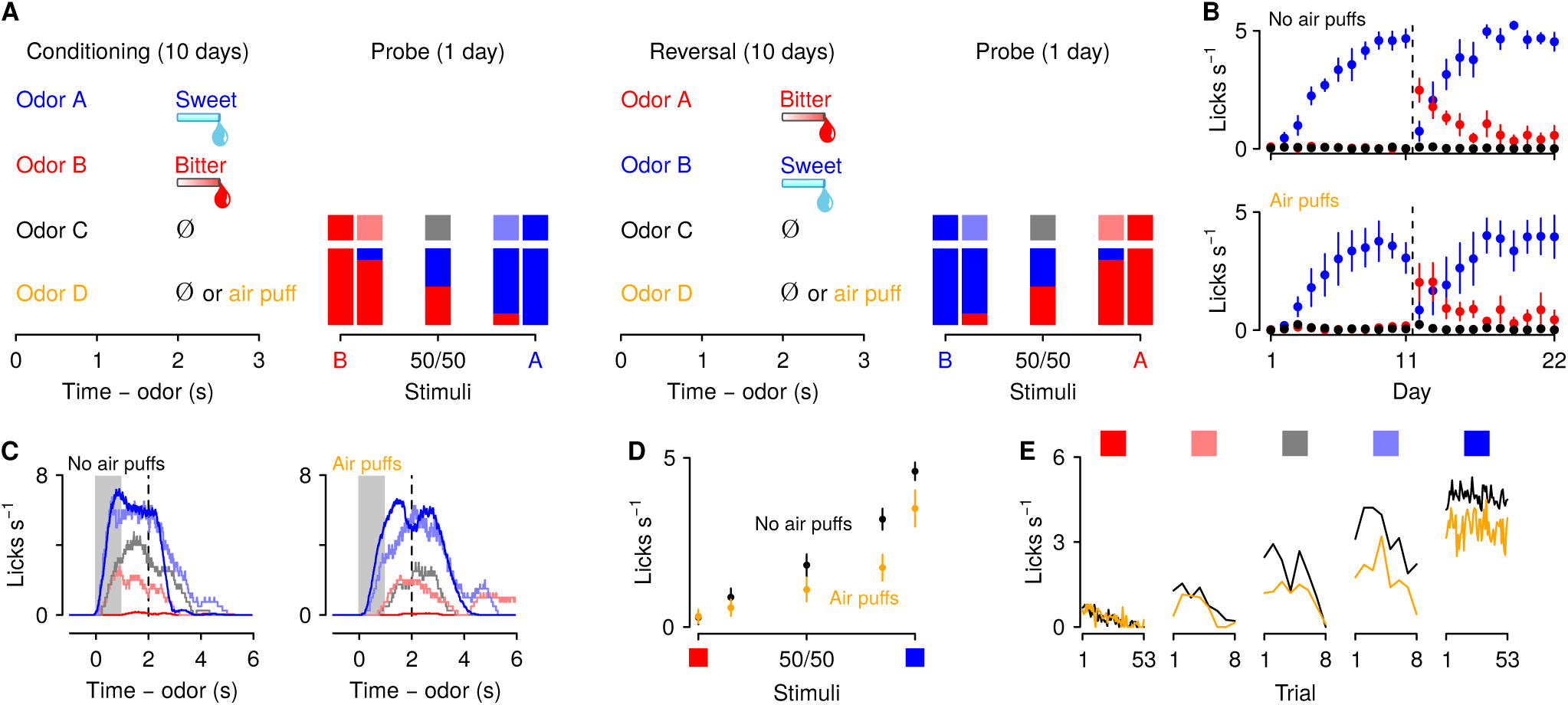
Behavioral responses to motivationally-significant predictive cues are modulated by punishment history. (A) Task design and experimental timeline. During ten conditioning sessions, odors (A, B, C and D) predicted an appetitive sucrose solution, an aversive denatonium solution, no outcome, and either no outcome or an unavoidable air puff, respectively. During the first probe test, A and B were mixed in three different ratios: 85%B/15%A (light red), 50%B/50%A (gray), 15%B/85%A (light blue). After completion of ten reversal training sessions, in which A and B contingencies were reversed, mice were re-trained in a second probe test. (B) Licking rates in no air puff (top, 7 mice) or air puff (bottom, 5 mice) groups across days, during odor and delay period, for sucrose (blue), denatonium (red), and no-outcome (black) trials. Dashed lines indicate reversals on day 11. (C) Licking behavior from a representative test session from a mouse without (left) and one with (right) exposure to air puffs. Color gradations between blue and red indicate odor mixtures as in (A). Gray bars indicate a period of odor presentation. Dashed lines indicate outcome delivery. (D) Licking rates during sucrose (blue square) and denatonium (red square) trials and during the eight probe trials for no air puff (black) and air puff (orange) groups, during odor and delay period. (E) Trial-by-trial licking rates during sucrose (blue square) and denatonium (red square) trials and during the eight probe trials for each ambiguous cue (light red, gray, light blue squares) for no air puff (black) and air puff (orange) groups, during odor and delay period. Line and error bars represent mean *±* SEM.

As predicted, mice in both air puff and no air puff groups quickly learned the CS-US associations: they showed increases in anticipatory licking responses to the positive, sucrose-predicting cue and in the delay before sucrose arrived across conditioning sessions, while withholding licking after sampling the negative, denatoniumpredicting cue (Figure 1B). Accordingly, a 3-factor ANOVA (session *×* cue *×* group) comparing licking behavior during sucrose and denatonium cue presentation and delay period demonstrated a significant main effect of session (*F*_1,9_ = 29.74, *p <* 0.01) and cue (*F*_1,1_ = 64.77, *p <* 0.01). Moreover, mice exposed to air puffs responded to the sucrose-predicting odor with fewer licks (cue *×* group interaction, *F*_1,1_ = 7.84, *p <* 0.05). During reversal learning, in which A and B contingencies were reversed, mice in both no air puff and air puff groups quickly re-adapted to the new associations (Figure 1B). Accordingly, a 3-factor ANOVA (session *×* cue *×* group) comparing licking rates during sucrose and denatonium cue presentation and delay period demonstrated a significant interaction between cue and session (*F*_1,9_ = 8.4, *p <* 0.01).

At the end of conditioning and reversal training, mice received a single probe test session. Behavioral responses from the initial probe test were not statistically different from data gathered in the second test and thus these sessions were analyzed together in the text. Licking rates for odor mixtures scaled with the proportion of the mixture that was the sucrose-predicting odor. This indicates that mice responded to parametrically varying ambiguous stimuli with smoothly varying behavioral responses (Figure 1C). Interestingly, mice exposed to air puffs responded to ambiguous odor mixtures with fewer licks during the anticipatory odor and delay period indicating that exposure to punishments biased decisions about ambiguous outcomes toward avoidance and away from approach (Figure 1D). Accordingly, a 2-factor ANOVA (cue *×* group) comparing licking rates during cue presentation and delay period in test days demonstrated a significant interaction between cue and group (*F*_1,4_ = 5.32, *p <* 0.01). Notably, the reduced licking to ambiguous cues in the air puff group was evident on the first trial of the probe test and persisted throughout all probe test trials (Figure 1E). Thus, the decline in responding was not due to effects of extinction in the probe test. Indeed, both groups showed similar extinction of responding to ambiguous cues across trials resulting from outcome omission. Accordingly, a three-factor ANOVA (cue *×* trial *×* group) revealed a significant interaction between cue and trial (*F*_1,14_ = 5.33, *p <* 0.01). Importantly, the interaction between cue, group and trial was not significant (*F*_1,14_ = 1.21, *p* = 0.27).

### Punishment history modulates mPFC neuronal responses to motivationally-relevant stimuli

Medial PFC (mPFC; also referred to as prelimbic cortex) is known to be involved in learning (Holland and Gallagher, 2004; Luk and Wallis, 2009; Alexander and Brown, 2011; Del Arco et al., 2017; Otis et al., 2017; Orsini et al., 2018), stress (Wellman, 2001; Cook and Wellman, 2004; Radley et al., 2004, 2005; Liston et al., 2006; Radley et al., 2006; Cerqueira et al., 2007; Wei et al., 2007; Liu and Aghajanian, 2008; Radley et al., 2008; Goldwater et al., 2009; Yuen et al., 2012; Adhikari et al., 2015), and uncertainty (Ernst and Paulus, 2005; Opris and Bruce, 2005; Sugrue et al., 2005; Bach et al., 2009; Levy et al., 2010; Orsini et al., 2018). These functions are critical for making predictions about previously unobserved stimuli. Such predictions derive from prior knowledge, as well as experience with the context of an environment. We thus asked whether mPFC neurons responded differently to motivationally-relevant stimuli in the presence or absence of punishment history.

We recorded action potentials extracellularly from 2,208 mPFC neurons in 12 mice, 5 exposed to air puffs (929 neurons), 7 unexposed (1,279 neurons), while mice performed the go/no-go discrimination task (Supplementary Fig. 1). Most neurons showed firing rate increases or decreases following odor cues, largely with persistent activity that lasted beyond cue offset. In particular, to characterize the responses of the population, we measured the temporal response profile of each neuron during sucrose trials by quantifying firing rate changes from denatonium trials in 100-ms bins using a receiver operating characteristic (ROC) analysis (Figure 2A). We calculated the area under the ROC curve (auROC), comparing sucrose trials to denatonium trials. This analysis determines to what extent an ideal observer could discriminate between activities on the two trial types. Values of 0.5 indicate no discriminability, whereas values of 0 or 1 indicate perfect discriminability. A large fraction of neurons (270 of 929 in mice exposed to air puffs and 325 of 1,279 in mice unexposed) showed auROC values greater than 0.7, indicating substantially greater activity for sucrose compared to denatonium trials, or values less than 0.3, indicating substantially greater activity for denatonium compared to sucrose trials (Figure 2A). These patterns were observed in mice exposed and unexposed to air puffs (Figure 2B). Neurons from mice exposed to air puffs responded robustly to both air puff-predicting cues and the air puffs themselves (Figure 2C; Wilcoxon rank sum test, *p <* 0.0001).

**Fig. 2.**
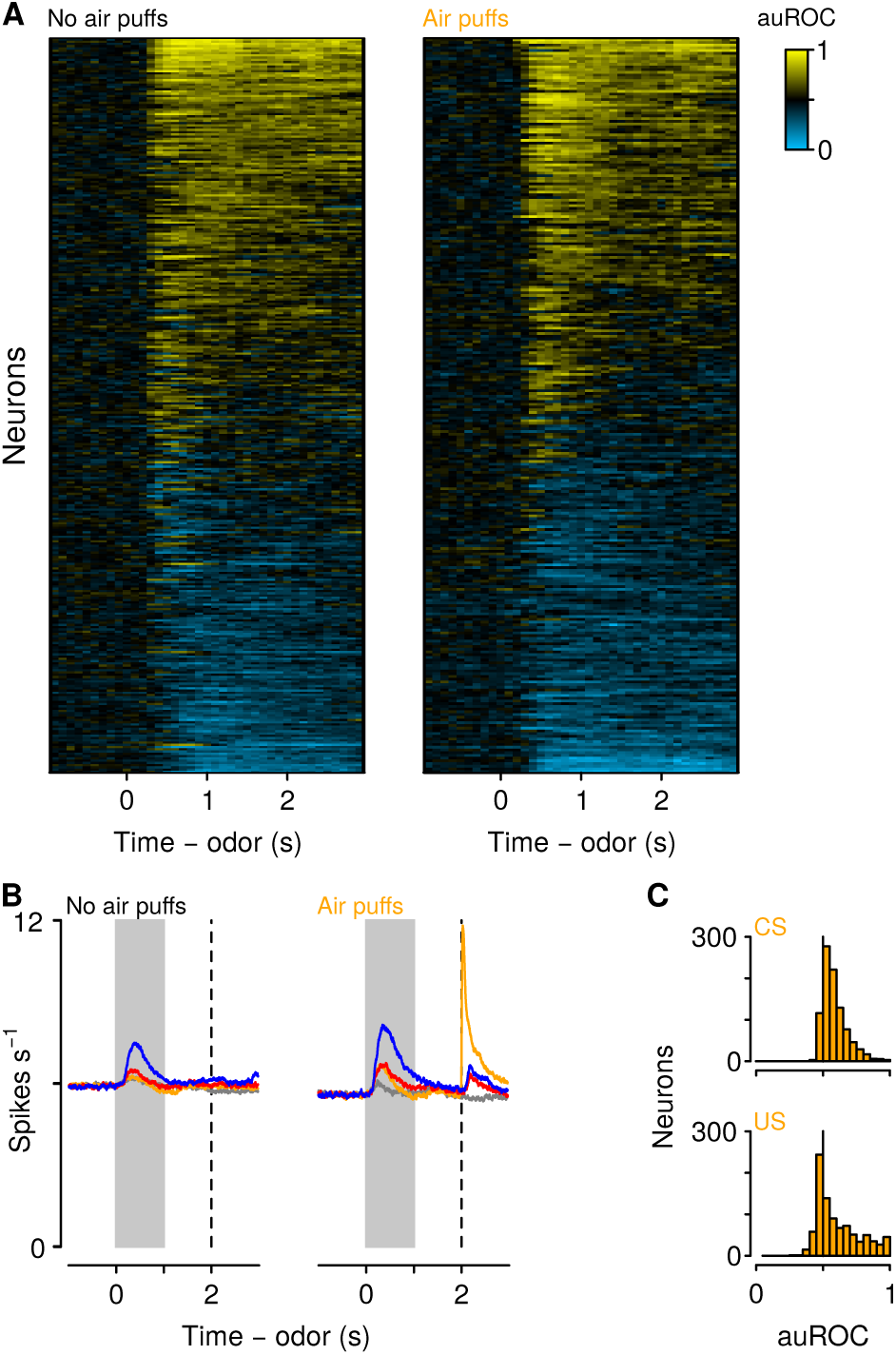
Neuronal responses across learning. (A) Discriminability (auROC) between sucrose and denatonium trials of all neurons in mice unexposed (left) and exposed (right) to air puffs. Increases (yellow) and decreases (cyan) in firing rate in sucrose trials relative to denatonium trials. Each row represents one neuron. (B) Average firing rates of all neurons with auROC values greater than 0.7 or less than 0.3 in at least one bin in no air puff (left) and air puff (right) mice during sucrose (blue), denatonium (red), no-outcome (gray) and air puff (orange) trials. Gray bars indicate a period of odor presentation. Dashed lines indicate outcome delivery. (C) auROC values for responses to air puff-predicting CSs (top) and air puff (bottom).

We next asked whether a history of air puffs changed the responses of mPFC neurons to aversion- and reward-predictive cues together with ambiguous stimuli. We recorded from 136 neurons from mPFC in the no air puff group and 106 neurons from mPFC in air puff exposed mice during the probe sessions (Figure 3). These populations included 37 in the no air puff group and 25 in the air puff trained group that exhibited auROC discrimination between sucrose and denatonium trials greater than 0.7 or less than 0.3 during the probe sessions. To compare firing rates of those neurons between mice exposed or unexposed to air puffs, we calculated a generalized linear model (Poisson regression) to predict spike counts during the cues as a function of air puff exposure and odor type. Cue-evoked firing rates were significantly higher in mice exposed to air puffs (Figures 3C and 3D; odor mixture *z* = 0.18 *±* 0.015, airpuff group *z* = 0.043 *±* 0.014, stimulus-group interaction *z* = 0.033 *±* 0.019, *p <* 0.001).

**Fig. 3.**
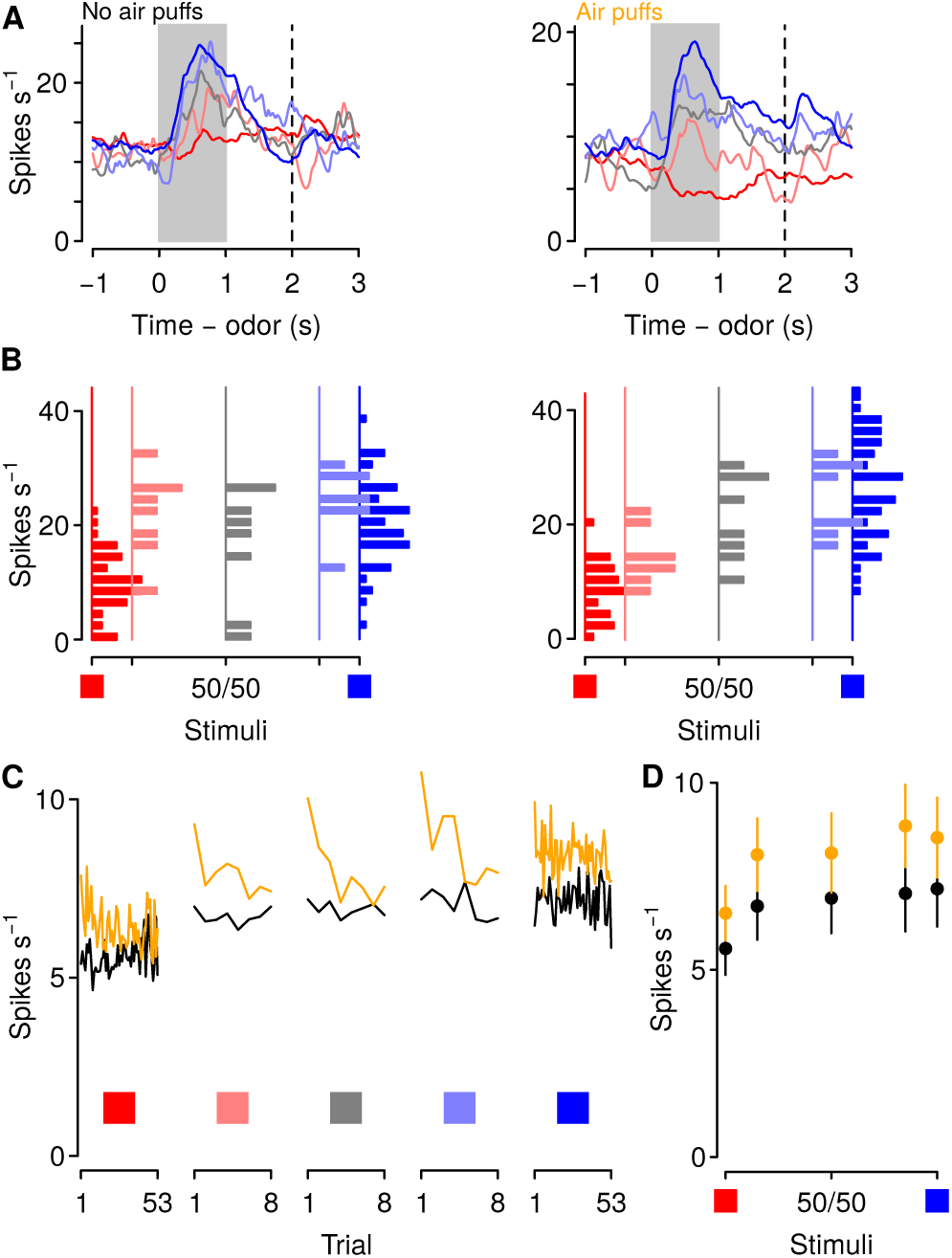
Punishment history increases mPFC firing rates to motivationally-relevant cues. (A) Average firing rates from example neurons in a mouse unexposed (left) and exposed (right) to air puffs. Red: denatonium trials. Blue: sucrose trials. Graded colors indicate mixtures as in Fig. 1A. Gray bars indicate a period of odor presentation. Dashed lines indicate outcome delivery. (B) Firing rates for the two example neurons in Fig. 3A during odor and delay period. (C) Trial-by-trial firing rates during sucrose (blue square) and denatonium (red square) trials and during the eight probe trials for each ambiguous cue (light red, gray, light blue squares) for no air puff (black) and air puff (orange) groups, during odor and delay period. (D) Mean *±* SEM firing rates during sucrose (blue square) and denatonium (red square) trials and during the eight probe trials for no air puff (black) and air puff (orange) groups, during odor and delay period.

### Punishment history modulates corticothalamic neuronal responses to motivationally-significant stimuli

The neural data described above suggests that a subgroup of mPFC neurons is modulated by punishment history. One of the major projection targets of the mPFC is the paraventricular nucleus of the thalamus (mPFC→PVT). Studies examining the role of PVT in regulating behavioral responses have found that PVT neurons are activated by cues or contexts previously associated with positive or negative emotional outcomes (Beck and Fibiger, 1995; Schiltz et al., 2007; Yasoshima et al., 2007; Choi et al., 2010; Igelstrom et al., 2010; Haight and Flagel, 2014; Hsu et al., 2014; Do-Monte et al., 2015; Kirouac, 2015; Matzeu et al., 2015; Penzo et al., 2015; Li et al., 2016; Zhu et al., 2016; Do-Monte et al., 2017). Additionally, activity in corticothalamic neurons suppresses both the acquisition and expression of conditioned reward seeking Otis et al. (2017). Thus, we hypothesized that mPFC→PVT neurons encode information about appetitive and aversive stimuli, and that this information is critical for weighing prior experience in those predictions.

To test this hypothesis, we performed projection-specific electrophysiological recordings from mPFC→PVT neurons while mice performed the go/no-go task. We expressed the light-gated ion channel channelrhodopsin-2 (ChR2, using adeno-associated viruses, AAV1-CaMKIIa-ChR2-eYFP) in pyramidal neurons of the mPFC and we implanted an optic fiber above the PVT, to activate mPFC→PVT cells antidromically (Figure 4A, B). Virus expression and optic fiber implantation were verified histologically (Supplementary Figure 2). At the end of each recording session, we used ChR2 excitation to observe antidromically-evoked spikes. For each neuron, we measured the response to light stimulation and the shape of spontaneous spikes (Figure 4C). To unequivocally identify mPFC→PVT neurons, ChR2-expressing cells in the mPFC were identified with axonal photostimulation and extracellular recordings in mPFC using a collision test (Figure 4D; Paintal, 1959; Bishop et al., 1962; Darian-Smith et al., 1963). Based on the parameters of cells that passed the collision test, we only included units that responded to light with a latency less than 15 ms and spiked in response to at least 70% of all pulses (in response to 10 Hz pulses; Figure 4D). These criteria are comparable to the fastest responses seen using antidromic stimulation with collision tests in corticothalamic neurons in sensory regions of neocortex (Swadlow and Weyand, 1981; Swadlow, 1998; Stoelzelet al., 2017).

**Fig. 4.**
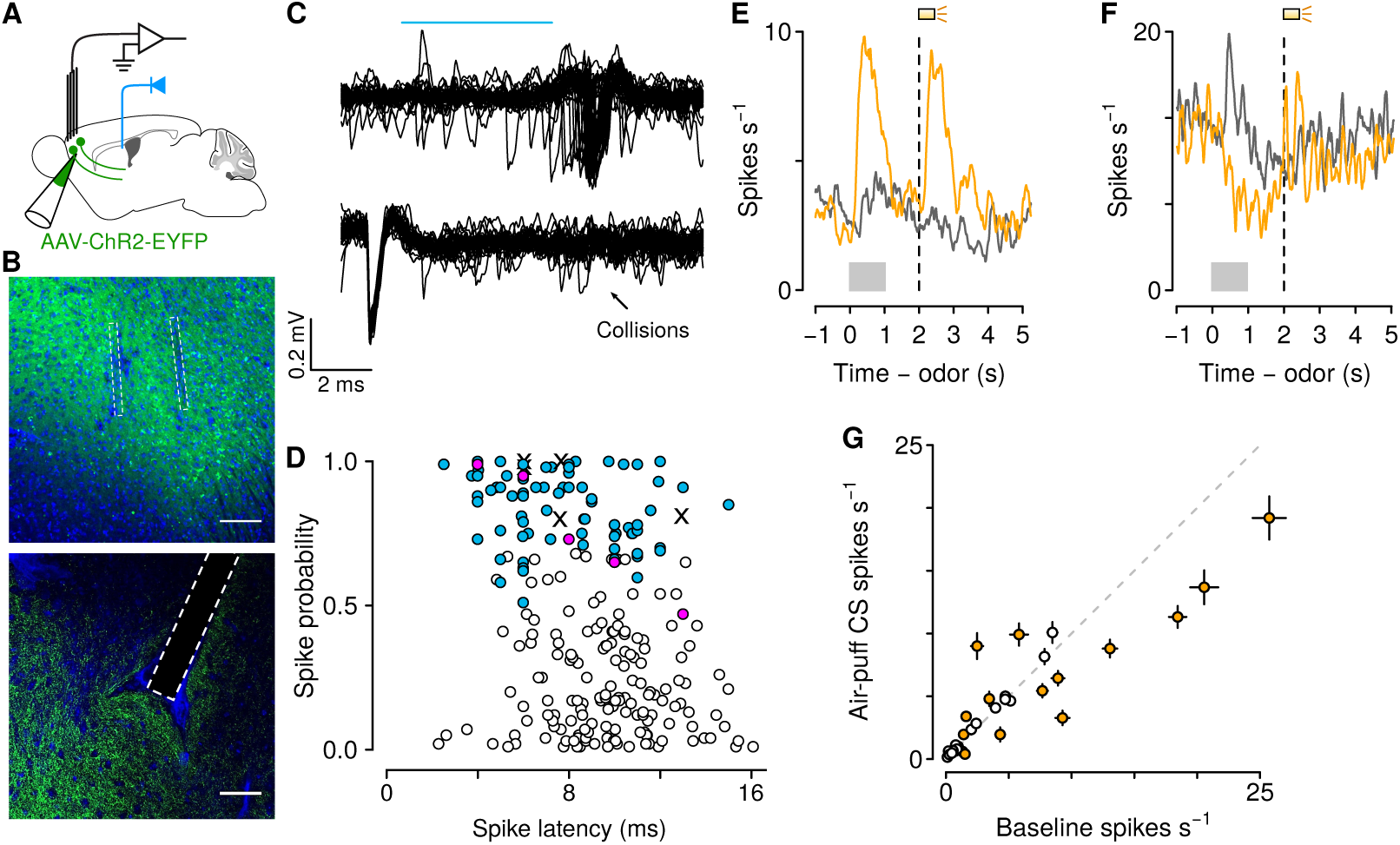
Air puff-predicting stimuli modulate mPFC→PVT neuron firing rates. (A) Schematic drawings of viral stereotaxic injection of AAV1-CaMKII-ChR2-eYFP and tetrode bundle into mPFC and optic fiber over PVT. (B) eYFP (green), and DAPI (blue) in mPFC (top) and PVT (bottom) coronal sections from BL6 mice that received AAV1-CaMKII-ChR2-eYFP and tetrode bundle into mPFC and an optic fiber over PVT (scale bar, 100 *µ*m). (C) Example of an identified corticothalamic neuron responding to a sequence of light stimuli (cyan) with action potentials (top) but not when the light stimuli followed spontaneous action potentials (bottom). (D) Antidromically-tagged corticothalamic neurons (blue) and antidromically-tagged corticothalamic neurons that passed collision tests (magenta). White points are neurons that were not identified. Crosses are neurons that passed collision tests, but were not recorded during behavior. (E-F) Average firing rates from example mPFC→PVT neurons showing firing rate increase (E) or decrease (F) to the air puff-predicting cue. Orange: air puff trials. Gray: CS*−* trials. Gray bars indicate a period of odor presentation. Dashed lines indicate outcome delivery. (G) Scatter plot showing relationship between the change in neural activity to the air puff-predicting cue compared to baseline firing activity. Orange: neurons in which the firing rate during the air puff-predicting cue was significantly different from baseline firing activity (*t*-test, *p <* 0.05).

We identified 39 and 45 neurons as projecting to PVT in mice unexposed and exposed to air puffs, respectively (Figure 4D; no air puff group: 18 and 21 neurons during conditioning and probe sessions, respectively; air puff group: 35 and 10 neurons during conditioning and probe sessions, respectively). We first asked whether these neurons responded to aversive stimuli. Previous studies have emphasized the role of the PVT in shaping behavioral responses to stimuli that predict aversive outcomes (Beck and Fibiger, 1995; Yasoshima et al., 2007; Hsu et al., 2014; Do-Monte et al., 2015; Penzo et al., 2015), but it’s unknown which of its inputs may drive those responses. We found that 43% of mPFC→PVT neurons recorded from mice in the air puff group showed firing rate changes (increases or decreases) in response to air-puff-predicting stimuli (Figures 4E-G). This demonstrates that mPFC→PVT neurons, thought to be involved in behavioral responses to aversive-predicting stimuli, are indeed modulated by those stimuli.

We next compared the responses of mPFC→PVT neurons to sucrose- and denatonium-predictive cues and ambiguous stimuli. In both groups, firing rates of mPFC→PVT neurons for odor mixtures scaled with the proportion of the mixture that was the sucrose-predicting odor, indicating that corticothalamic neurons responded to parametrically varying ambiguous stimuli with smoothly varying neural responses (Figure 5A-B). Interestingly, corticothalamic neurons from mice exposed to air puffs responded to the sucrose-predictive cue and ambiguous odor mixtures with higher phasic activity during the anticipatory odor and delay period, indicating that exposure to punishments biases neural responses of corticothalamic neurons to reward-predictive and ambiguous stimuli (Figure 5C-D; Poisson regression, odor mixture *z* = 0.26 *±* 0.10, *p <* 0.01, air-puff group *z* = 0.73 *±* 0.15, *p <* 0.01, stimulus-group interaction *z* = 0.27 *±* 0.22, *p >* 0.2).

**Fig. 5.**
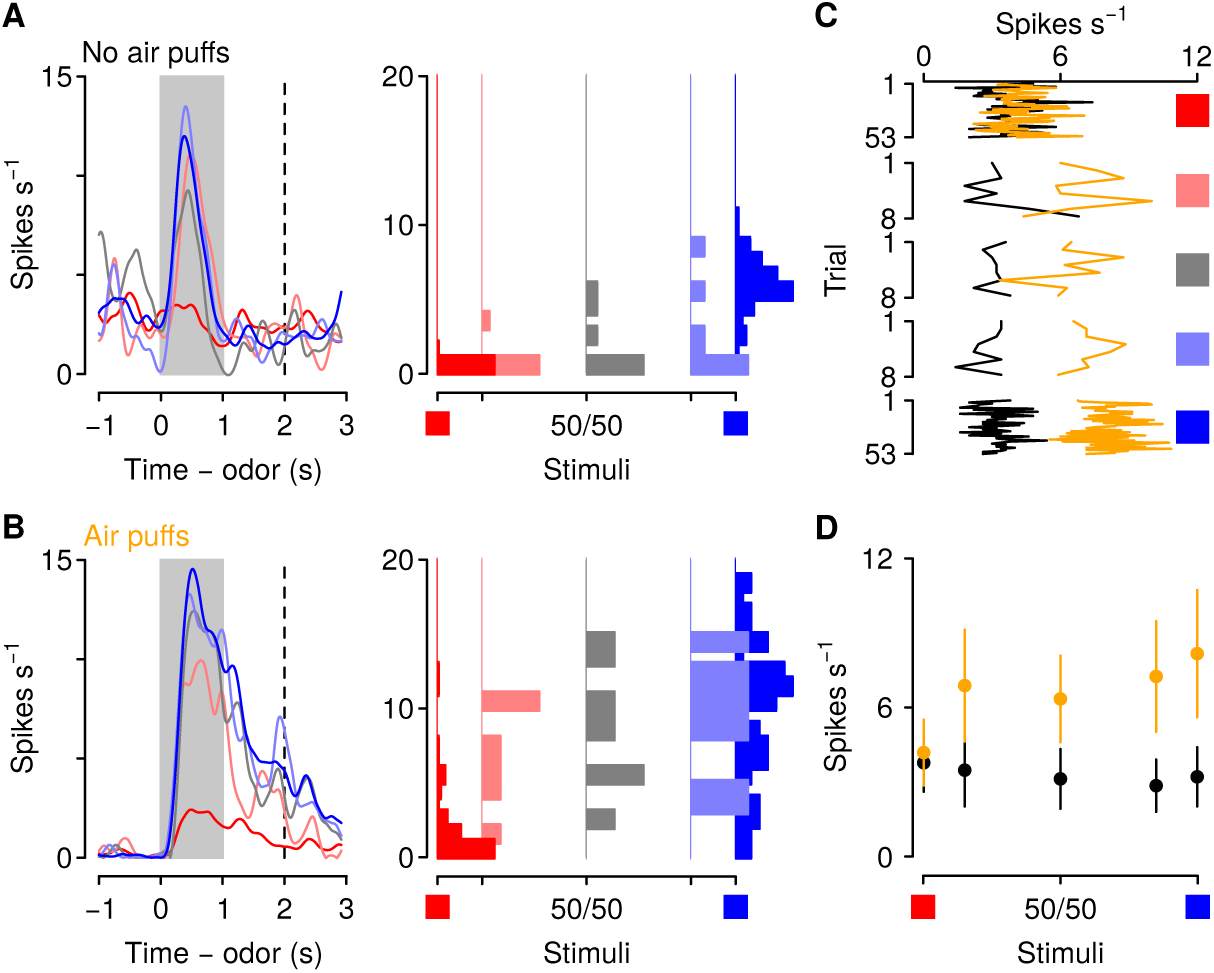
Punishment history increases corticothalamic firing rates to motivationally-relevant cues. (A-B) Average firing rates (left) and firing rates during odor and delay period (right) from example neurons in a mouse unexposed (A) and exposed (B) to air puffs. Red: denatonium trials. Blue: sucrose trials. Graded colors indicate mixtures as in Fig. 1A. Gray bars indicate a period of odor presentation. Dashed lines indicate outcome delivery. (C) Trial-by-trial firing rates during sucrose (blue square) and denatonium (red square) trials and during the eight probe trials for each ambiguous cue (light red, gray, light blue squares) for no air puff (black) and air puff (orange) groups, during odor and delay period. (D) Mean *±* SEM firing rates during sucrose (blue square) and denatonium (red square) trials and during the eight probe trials for no air puff (black) and air puff (orange) groups, during odor and delay period.

### Excitation of corticothalamic neurons modulates behavioral responses to motivationally-relevant cues

The neural data described above suggests that elevated activity in corticothalamic neurons to the aversion- and reward-predictive cues together with the ambiguous cues is critical for modulating behavioral response to those stimuli. To provide a more specific test of this hypothesis, we next used optogenetic methods to activate corticothalamic neurons selectively at the time of presentation of those cues in mice trained in the same go/no-go task described above but with no exposure to air puffs. Mice received bilateral infusions of either AAV1/5-CaMKIIa-ChR2-eYFP or AAV1/5-CaMKIIa-eYFP (control) into mPFC; expression was verified histologically (Supplementary Figure 3). Mice also received fiber optic implants over PVT (Figure 6A). Three weeks after surgery, these mice began training in the go/no-go discrimination task and, during the probe session, light was delivered into the PVT in 5 random trials of all sucrose and denatonium trials and in half of all the ambiguous trials, in the cue and delay period (Figure 6B). We used a 10 Hz frequency stimulation because it resembled the firing rate of corticothalamic neurons in response to ambiguous stimuli observed in neural recordings; we also used a 20 Hz frequency in order to get a frequency response curve. All mice underwent conditioning sessions and there were neither main effects nor any interactions of group on conditioned responding across conditioning (*F <* 1.14; *p >* 0.33) (Figure 6C). During the subsequent probe test, ChR2 mice showed a reduction in response to denatonium- and sucrose-predictive cues and ambiguous cues in the trials in which light was delivered in a frequency-dependent manner, whereas eYFP mice that received the same treatment responded equally in trials with or without light delivery (Figure 6D-E). Accordingly, a 3-factor ANOVA (cue *×* stimulation *×* group) comparing licking behavior during cue presentation and delay period in stimulated versus unstimulated trials in eYFP, 10 Hz and 20 Hz groups revealed a significant main effect of cue (*F*_1,4_ = 13.8, *p <* 0.01) and stimulation (*F*_1,1_ = 30.11, *p <* 0.01). Moreover, there was a significant interaction between cue and group (*F*_1,8_ = 4.8, *p <* 0.01) and stimulation and group (*F*_1,2_ = 4.68, *p <* 0.05).

**Fig. 6.**
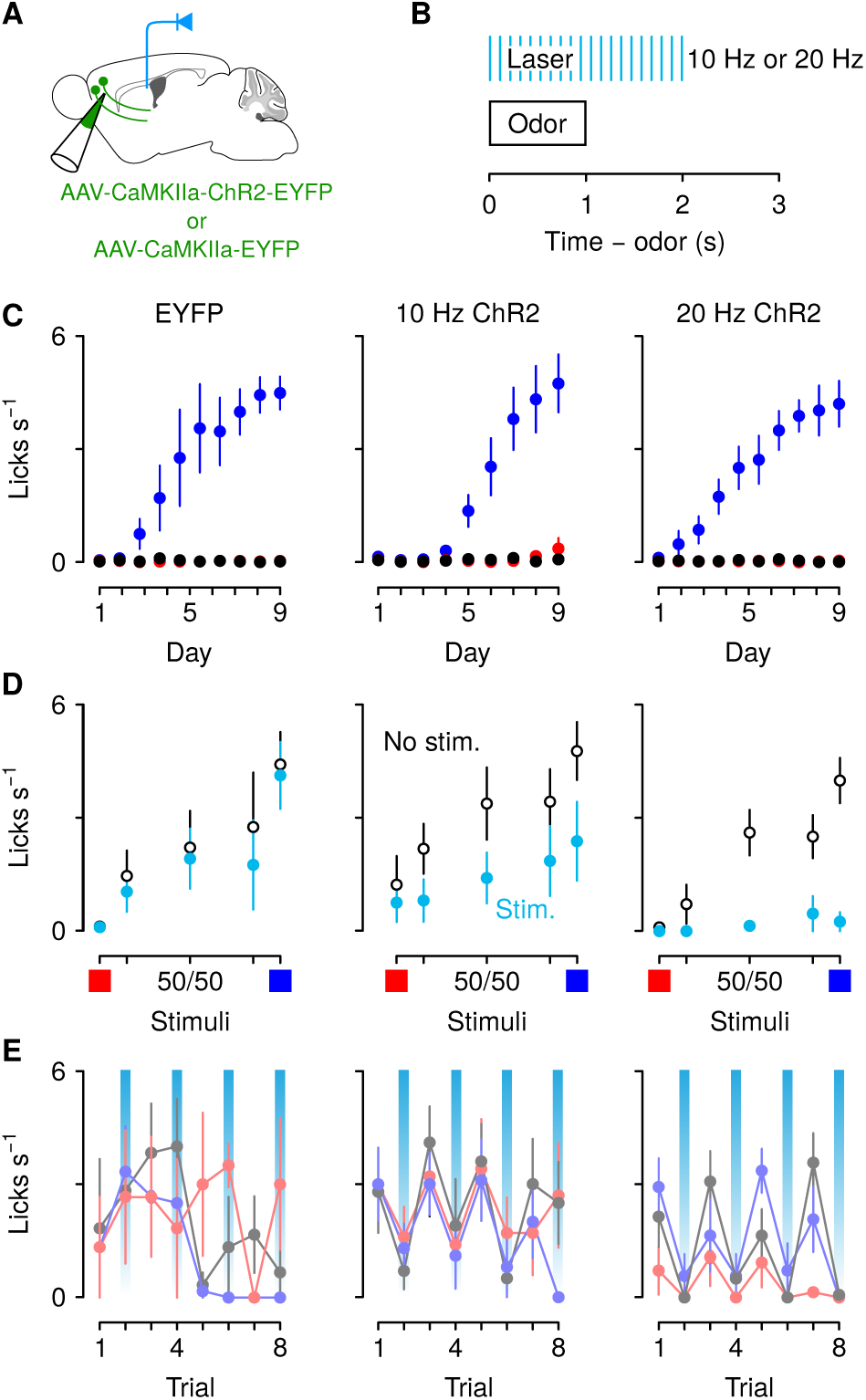
Optogenetic excitation of corticothalamic neurons negatively biases responses to motivationally-relevant stimuli. (A) Schematic of viral stereotaxic injection of AAV1/5CaMKIIa-ChR2-eYFP or AAV1/5-CaMKIIa-eYFP into mPFC and optic fiber over PVT. (B) Optical stimulation was delivered during presentation of the cue and during the 1 s delay before outcome delivery. (C) Licking rates in eYFP (left, *n* = 3), 10 Hz ChR2-eYFP (center, *n* = 5), and 20 Hz ChR2-eYFP (right, *n* = 7) groups across conditioning, during odor and delay period, for sucrose (blue), denatonium (red), and no-outcome (black) trials. (D) Licking rates during sucrose (red square) and denatonium (blue square) trials and during the eight probe trials (light blue, gray and light red squares) for eYFP (left), 10 Hz ChR2eYFP (center) and 20 Hz ChR2-eYFP (right) groups, during odor and delay period, with (light blue) or without (white) laser stimulation. (E) Trial-by-trial licking rates during 85%A/15%B (light red), 50%A/50%B (gray), 15%A/85%B (light blue) trials for eYFP (left), 10 Hz ChR2-eYFP (center) and 20 Hz ChR2-eYFP (right) groups, during odor and delay period, with (light blue shadows) or without laser stimulation. Line and error bars represent the mean *±* SEM.

Stimulation of the same neurons did not, however, disrupt licking for an unpredictable reward. Outside of the task, we delivered randomly-timed sucrose rewards (3 *µ*l). Mice licked at high rates to consume unexpected rewards (Supplementary Figure 4). Importantly, stimulation of mPFC→PVT axons during reward delivery at the same frequencies used in the behavioral task in half of the total number of trials did not change the number of licks in response to these rewards (Supplementary Figure 4). Accordingly, a 2-factor ANOVA (frequency of stimulation *×* group) comparing licking behavior during stimulated and unstimulated trials revealed no significant main effect or interaction with group (all *F <* 0.23, *p >* 0.65). Thus, uncued licking is not altered by optogenetic excitation of mPFC→PVT cells and the optogenetic effects are not due to a light-induced impairment in licking in general.

## Discussion

In this study, we examined the projection from mPFC to PVT in mice approaching or avoiding negative and positive valence-predictive stimuli. We found that a history of punishments negatively biased behavioral responses to those motivationally-relevant stimuli while selectively increasing excitatory responses of PVT-projecting mPFC neurons. Indeed, mice exposed to punishments showed reduced approach behavior to reward-predictive and ambiguous stimuli than mice unexposed to punishments. Moreover, artificially increasing activity from mPFC to PVT quantitatively mimicked the punishment-induced negative behavioral bias.

Cognitive processes—appraisals of stimuli, events and situations—play an important role in generating affective states, and vice versa, these affective states influence cognitive functioning by inducing attentional, memory, and judgment biases (Lerner and Keltner, 2000; Haselton et al., 2009; Harding et al., 2004; Enkel et al., 2010; Rygula et al., 2012; Papciak et al., 2013; Rygula et al., 2013; Parker et al., 2014; Rygula et al., 2014). Among brain regions that are engaged in affective processing, the mPFC has long been implicated in adaptive responding by signaling information about expected outcome and by regulating sensitivity to reward and punishment (Holland and Gallagher, 2004; Luk and Wallis, 2009; Alexander and Brown, 2011; Del Arco et al., 2017; Orsini et al., 2018). Here, we found that firing rates in mPFC neurons reflect cue-evoked expectations for aversion- and reward-predictive cues and were enhanced in mice exposed to a mild punishment, which correlates with a reduction in anticipatory responses to the same stimuli. Those findings are consistent with a negative bias and suggests that negative events can bias decisions by altering the activity of mPFC neurons. Several studies have implicated the prefrontal network in the pathophysiology of affective disorders (Phillips et al., 2003; Drevets et al., 2008) and chronic stress—a crucial factor in increasing the risk of developing affective disorders— has profound detrimental effects on the anatomy and physiology of mPFC neurons (Wellman, 2001; Cook and Wellman, 2004; Radley et al., 2004, 2005; Liston et al., 2006; Radley et al., 2006; Cerqueira et al., 2007; Wei et al., 2007; Liu and Aghajanian, 2008; Radley et al., 2008; Goldwater et al., 2009; Yuen et al., 2012; Adhikari et al., 2015). In particular, chronic stress induces significant regression of the apical dendrites of pyramidal neurons in mPFC (Cook and Wellman, 2004; Radley et al., 2004; Liston et al., 2006; Goldwater et al., 2009), which may in turn impact mPFC function.

The mPFC densely projects to subcortical motivationally relevant processing structures, including PVT, amygdala, hippocampus and nucleus accumbens (Vertes, 2004; Li and Kirouac, 2012). Among all brain regions that receive strong projections from the mPFC, PVT has long been considered a stress detector and implicated in the emergence of adaptive responding to stress (Chastrette et al., 1991; Sharp et al., 1991; Cullinan et al., 1995; Bubser and Deutch, 1999; Spencer et al., 2004; Hsu et al., 2014; Do-Monte et al., 2015; Penzo et al., 2015; Zhu et al., 2016; Do-Monte et al., 2017; Beas et al., 2018; Choi et al., 2019). On the other hand, PVT has also been considered a potential mediator of motivated behavior responding to both food- and drug-associated cues (Schiltz et al., 2005; Igelstrom et al., 2010; Martin-Fardon and Boutrel, 2012; James and Dayas, 2013; Browning et al., 2014; Haight and Flagel, 2014; Li et al., 2016). Consistent with these findings that put PVT in a unique position to integrate information about positive and negative motivationally relevant cues and translate it into adaptive behavioral responses, we found that mPFC neurons projecting to the PVT maintain cue-evoked expectations for motivationally-relevant outcomes and their neural activity was enhanced in mice exposed to punishments. Moreover, by mimicking this punishment-induced increase of PVT-projecting mPFC neuronal activity with a selectively optogenetic activation of this pathway, we observed a reduction in anticipatory responses to the predictive cues. These findings suggest that information about cue interpretation is transferred from mPFC to PVT and this pathway is crucial for an adaptive responding toward those stimuli as a function of previous experiences. These results are also consistent with a recent study showing that activity in mPFC neurons projecting to the PVT suppresses both the acquisition and expression of conditioned reward seeking (Otis et al., 2017). Based on recent evidence showing two genetically, anatomically and functionally distinct cell types across the anteroposterior axis of the PVT (Gao et al., 2020), it will be interesting to investigate in future studies whether the information from mPFC is transferred to anatomically or molecularly segregated cell types or projection-specific neurons within the PVT.

Understanding the underlying mechanisms by which information of cue interpretation is updated as a function of prior experience in the mPFC→PVT circuit is crucial for delving deeper into the brain’s neuronal connectivity underlying cognitive bias behaviors. Importantly, proper tuning of this network has been shown to be exerted by robust neuromodulation from ascending catecholaminergic systems (Arnsten et al., 2012) and maladaptive processing of these systems has been implicated in cognitive deficits associated with several affective disorders (Enkel et al., 2010; Kukolja et al., 2008). For example, acute pharmacological stimulation of the serotonergic and dopaminergic systems has been shown to influence cognitive bias in rodents (Rygula et al., 2014). In particular, citalopram, a selective serotonin reuptake inhibitor, and amphetamine, a powerful psychostimulant, both induced a positive cognitive processing bias (Rygula et al., 2014). Those results are important from a clinical point of view, knowing that negative cognitive bias lies at the core of the pathophysiology of several affective disorders and it has been extensively studied in humans (Wright and Bower, 1992; MacLeod and Byrne, 1996; Beck, 2008). For example, it has been shown that patients with anxiety and depression interpret ambiguous information with a negative bias (Schwarz and Clore, 1983; Eysenck et al., 1987; Wright and Bower, 1992; MacLeod and Byrne, 1996; Lawson et al., 2002; Beck, 2008; Chan et al., 2008; Pizzagalli et al., 2008; Dearing and Gotlib, 2009). Thus, yielding a clearer vision of how cognitive biases develop and act in several chronic and debilitating neuropsychiatric disorders may offer an unprecedented opportunity for designing novel treatments, aimed at ameliorating the proper functional tuning and connectivity of prefrontal-orchestrated neuronal circuits.

In summary, by means of a mild punishment we induced a negative bias in mice which was linked to a hyperactivity in neural responses of PVT-projecting mPFC neurons. Artificial activation of the same pathway recapitulated the behavioral outcome. Thus, our results highlight a fundamental role for the mPFC→PVT circuit in shaping adaptive responses by modulating predictions about imminent motivationally-relevant outcomes as a function of prior experience.

## Materials and methods

### Subjects

Wild-type C57BL/6J male mice (The Jackson Laboratory, 000664), 8-10 weeks old at the time of surgery, were housed in a reverse 12-hour light-dark cycle room (lights on at 20:00). All mice were given ad libitum water except during testing periods. During behavioral testing, mice were water deprived by giving 1ml of water per day. Food was freely available throughout the experiments. All testing was conducted in accordance with the National Institutes of Health Guide for the Care and Use of Laboratory Animals and approved by the Johns Hopkins University Animal Care and Use Committee.

### Stereotaxic surgeries

All mice were surgically implanted with custom-made titanium head plates using dental adhesive (C&B-Metabond, Parkell) under isoflurane anesthesia (1.0-1.5% in O_2_). Surgeries were conducted under aseptic conditions and analgesia (ketoprofen, 5 mg/kg and buprenorphine, 0.05-0.1 mg/kg) was administered postoperatively. Mice recovered for 7-10 days before starting behavioral testing.

For electrophysiological experiments, we implanted unilaterally a custom microdrive containing 8 drivable tetrodes made from nichrome wire (PX000004, Sandvik) and positioned inside 39 ga polyimide guide tubes. We targeted mPFC under stereotaxic guidance at 2.2 mm anterior and 0.4 mm lateral to bregma and 1.6 mm ventral to the skull. Tetrodes were advanced subsequently into final positions in mPFC during recording. For identifying corticothalamic neurons, an optic fiber was implanted over PVT under stereotaxic guidance at 1.4 mm anterior and 1.3 mm lateral to bregma and 3.6 mm ventral to the skull with a 22.5*°* angle.

For optogenetic experiments, AAV1/5-CamKIIa-hChR2(H134R)-eYFP or AAV1/5-CamKIIa-eYFP (AAV5: from UNC GTC Vectore Core; AAV1: from Addgene) was injected bilaterally in mPFC under stereotaxic guidance at 2.2 mm anterior and 0.3 mm lateral to bregma and 1.6 mm ventral to the skull. pAAV-CaMKIIa-hChR2(H134R)-EYFP was a gift from Karl Deisseroth (Addgene viral prep 26969-AAV1; http://n2t.net/addgene:26969; RRID:Addgene 26969) (Lee et al., 2010). A total of 300 nl of virus (titer *∼* 10^13^ GC/mL) per hemisphere was delivered at the rate of 1 nl/s (MMO-220A, Narishige). The injection pipette was left in place for 5 min after each injection. Optic fibers (200 *µ*m diameter, 0.39 NA, Thorlabs) were implanted bilaterally over mPFC (at 2.2 mm anterior and 0.6 mm lateral to bregma and 1.3 mm ventral to the skull with a 10*°* angle) or unilaterally over PVT (at 1.4 mm anterior and 1.3 mm lateral to bregma and 3.6 mm ventral to the skull with a 22.5*°* angle).

### Behavioral task

Following recovery from surgery, water-restricted mice were habituated for 3 days while head-fixed before training on the go/no-go task. Each mouse performed behavioral tasks at the same time of day (between 08:00 a.m. and 2:00 p.m.). All behavioral tasks were performed in dark, sound-attenuated chamber, with white noise delivered between 2-60 kHz (L60 Ultrasound Speaker, Pettersson). Odors were delivered with a custom-made olfactometer (Cohen et al., 2012). Each odor was dissolved in mineral oil at 1:10 dilution. Diluted odors (30 *µ*l) were placed on filter-paper housing (Whatman, 2.7 *µ*m pore size). Odorized air was further diluted with filtered air by 1:10 to produce a 1.0 l/min flow rate. Licks were detected by charging a capacitor (MPR121QR2, Freescale). Task events were controlled with a microcontroller (ATmega16U2 or ATmega328). Reinforcements were 3 *µ*l of sucrose (an appetitive sweet solution), denatonium (an aversive bitter solution) or air puff (40 psi), delivered using solenoids (LHDA1233115H, The Lee Co). Intertrial intervals (ITIs) were drawn from an exponential distribution with a rate parameter of 0.3, with a maximum of 30 s. This resulted in a flat ITI hazard function, ensuring that expectation about the start of the next trial did not increase over time (Luce, 1986). The mean ITI was 7.2 s (range 2.4-30.0 s).

Mice underwent 10 conditioning sessions. In each session, mice received 50 1-s presentations of four different olfactory stimuli (A, B, C, and D). The order of odor presentations was randomized among mice and among sessions. For all conditioning, A, B, C, and D consisted of (+)-limonene, p-cymene, penthylacetate, and acetophenone, respectively (counterbalanced). 1-s after termination of A, sucrose was delivered and 1-s after termination of B, denatonium was delivered. C was paired with no reinforcement. In the air puff group, 1-s after termination of D, an unavoidable air puff was delivered to their right eye, while in the control group, D was paired with no reinforcement. 4-s after the presentation of each odor, a vacuum was activated to remove any residual of sucrose or denatonium. After the activation of the vacuum, there was a fixed 3 second delay and then the variable ITI will follow. After completion of conditioning training, mice received a single extinction probe session. During the probe session, the 4 conditioning odors were continued to be presented (53 trials for each odor), but mice also received eight non-reinforced presentations of three mixtures of varying proportions of A and B odors: 85%A/15%B, 50%A/50%B, 15%A/85%B. These odor mixture trials were interleaved with the 4 conditioning odor trials in a randomized order.

In mice designated for the mPFC electrophysiological experiments, following the probe test, mice underwent reversal learning, in which A and B were reversed. 1 s after termination of A, denatonium was delivered and 1 s after termination of B, sucrose was delivered. C and D were continued to be presented as in conditioning. Then, mice received another single extinction probe session, identical to the one received after conditioning. Neural data from the initial extinction days were not statistically different from data gathered in later rounds of training and thus these neurons were analyzed together in the text.

In mice designated for the PVT-projecting mPFC electrophysiological experiments, following the probe test, mice repeated three days of conditioning and then underwent additional rounds of probe test days in order to acquire additional data. This was done up to three times for a given mouse. Neural data from the initial extinction days were not statistically different from data gathered in later rounds of training and thus these neurons were analyzed together in the text.

In mice designated for the optogenetic experiments, training began approximately 3 weeks after viral injection and fiber implantation, and light (473 nm, 10-12 mW) was delivered into the PVT during the probe session. During the behavioral task, light was delivered in half of all the ambiguous trials, during the cue and delay epoch. Moreover, light was also delivered in 5 random trials of all sucrose and denatonium trials, during the cue and delay epoch. The primary measure of conditioning to cues was the number of licks during odor presentation and the second preceding reinforcement delivery. During the un-cued stimulation trials, light was delivered in half of all trials, during the presentation of reward delivery and lasted for 1500 ms.

### Electrophysiology

Throughout the discrimination task, mice were attached to the recording cable and before each session, tetrodes were screened for activity. Active tetrodes were selected for recording, and the session was begun. On the rare occasion that fewer than 4 of 8 tetrodes had single units, the tetrode assembly was advanced 40 or 80 *µ*m at the end of the session. Otherwise, the tetrode assembly was kept in the same position between sessions until the probe test day. After the extinction probe test, the tetrode assembly was advanced 80 *µ*m regardless of the number of active tetrodes in order to acquire activity from a new group of neurons in any subsequent training.

We recorded extracellularly (Digital Lynx 4SX, Neuralynx Inc.) from multiple neurons simultaneously at 32 kHz using custom-built screw-driven microdrives with 8 tetrodes coupled to a 200 *µ*m fiber optic (32 channels total). All tetrodes were gold-plated to an impedance of 200-300 kΩ prior to implantation. Spikes were bandpass filtered between 0.3-6 kHz and sorted online and offline using Spikesort 3D (Neuralynx Inc.) and custom software written in MATLAB. To measure isolation quality of individual units, we calculated the L-ratio (Schmitzer-Torbert et al., 2005) and fraction of interspike interval (ISI) violations within a 2 ms refractory period. All single units included in the dataset had an L-ratio less than 0.05 and less than 0.1% ISI violations. We only included units that had a firing rate of greater than 0.5 spikes s^*−*1^ over the course of the recording session.

### Optogenetic identification

To verify that our recordings targeted corticothalamic neurons, at the end of daily recording sessions, we used channelrhodpsin excitation to observe stimulation-locked spikes, by delivering 3-5 ms pulses of 473 nm light at 15 mW using a diode-pumped solid-state laser (Laserglow), together with a shutter (Uniblitz). Spike shape was measured using a broadband signal (0.1 Hz-9 kHz) sampled at 32 kHz. This ensured that particular features of the spike waveform were not missed. We delivered 10 trains of light (10 pulses per train, 10 s between trains) at 10 Hz, resulting in 100 total pulses. For collision tests, we delivered light over PVT (2 ms pulse, 473 nm, 15 mW) triggered by a spontaneous action potential. Identified neurons did not show stimulus-locked spikes following spontaneous spikes (“collisions”).

### Histology

At the end of behavioral testing, all mice were deeply anesthetized and then transcardially perfused with 4% paraformaldehyde (wt/vol). The brains were removed and processed for histology using standard techniques. For the electrophysiological experiments, we verified recording sites histologically with electrolytic lesions (15 s of 10 *µ*A direct current across two wires of the same tetrode). For optogenetic experiments, virus expression was examined using a confocal microscope (Zeiss LSM 800). After histological verification, mice with incorrect virus injection or tetrode implantation were excluded from data analysis.

### Data analysis

All analyses were performed with MATLAB (Mathworks) and R (http://www.rproject.org/). All data are presented as mean *±* SEM unless reported otherwise. All statistical tests were twosided. For nonparametric tests, the Wilcoxon rank-sum test was used, unless data were paired, in which case the Wilcoxon signed-rank was used. To estimate neuronal learning rates (Figure 2), we used logistic functions of the form 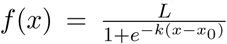. Learning rates are estimates of the *k* parameter.

## Acknowledgements

We thank T. Shelley for machining, and G. Schoenbaum, M. Pignatelli. and the Cohen Lab for comments. This work was supported by Life Sciences Research Foundation (F.L.), F30MH110084 (B.A.B.), Klingenstein-Simons, MQ, NARSAD, Whitehall, R01DA042038, and R01NS104834 (J.Y.C.), and P30NS050274.

## Author contributions

F.L., Z.S., A.J.C., and B.A.B. collected data. F.L. and J.Y.C. designed experiments, analyzed data, and wrote the paper.

## Author information

Correspondence should be addressed to J.Y.C. (http://jeremiah.cohen@jhmi.edu).

---

**Supplementary Figure 1.**
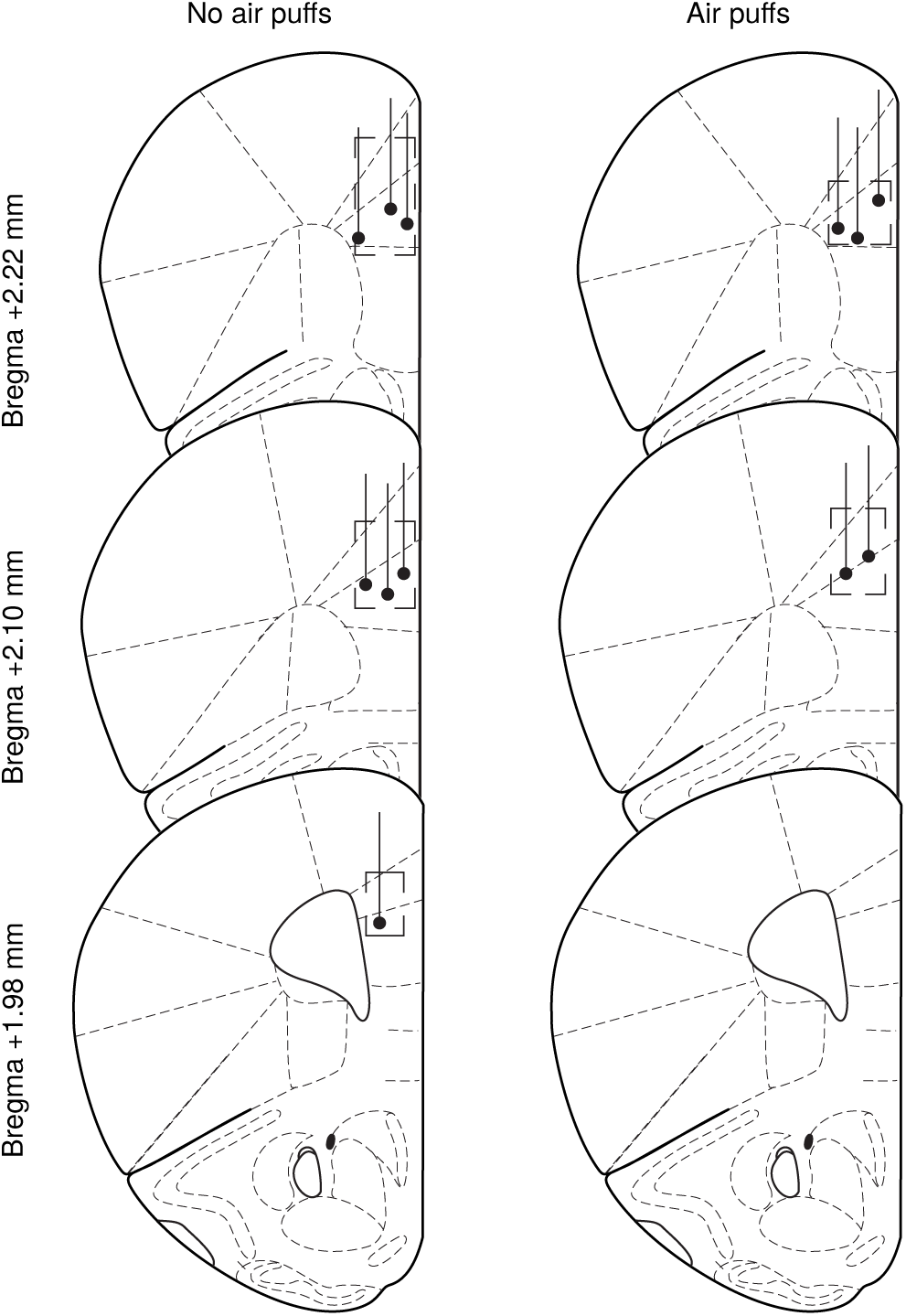
Drawings illustrate recording sites in mPFC in no air puff (left) and air puff exposed (right) mice. Boxes indicate approximate location of recording sites in each mouse, taking into account any vertical distance traveled during training and the approximate lateral spread of the tetrode bundle.

**Supplementary Figure 2.**
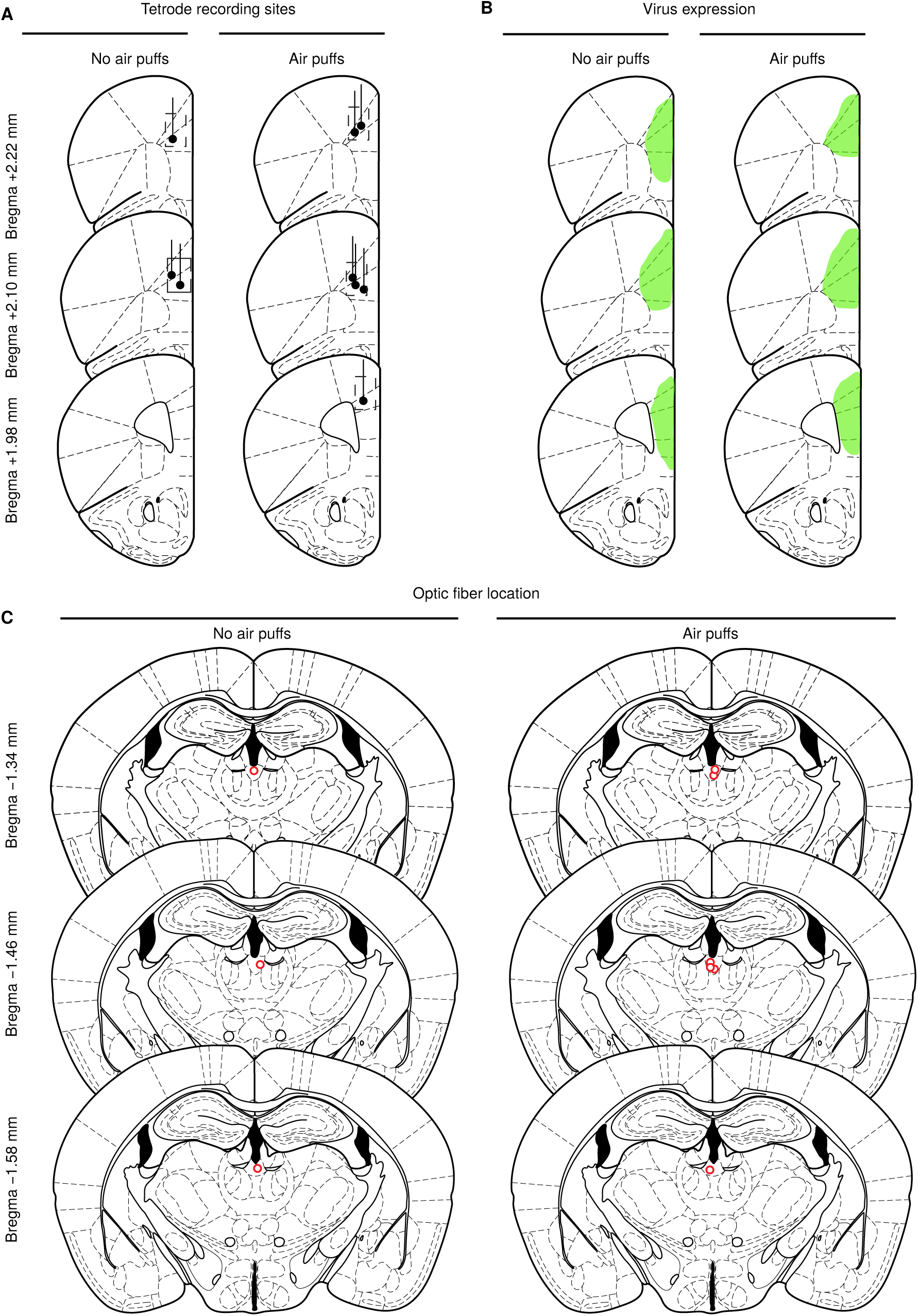
(A) Drawings illustrate recording sites in mPFC in no air puff (left, N=3) and air puff exposed (right, N=6) mice. Boxes indicate approximate location of recording sites in each mouse, taking into account any vertical distance traveled during training and the approximate lateral spread of the tetrode bundle. (B) Traces showing virus expression in no air puff (left) and air puff (right) groups. (C) Locations of fiber tips in no air puff (left) and air puff (right) groups in PVT.

**Supplementary Figure 3.**
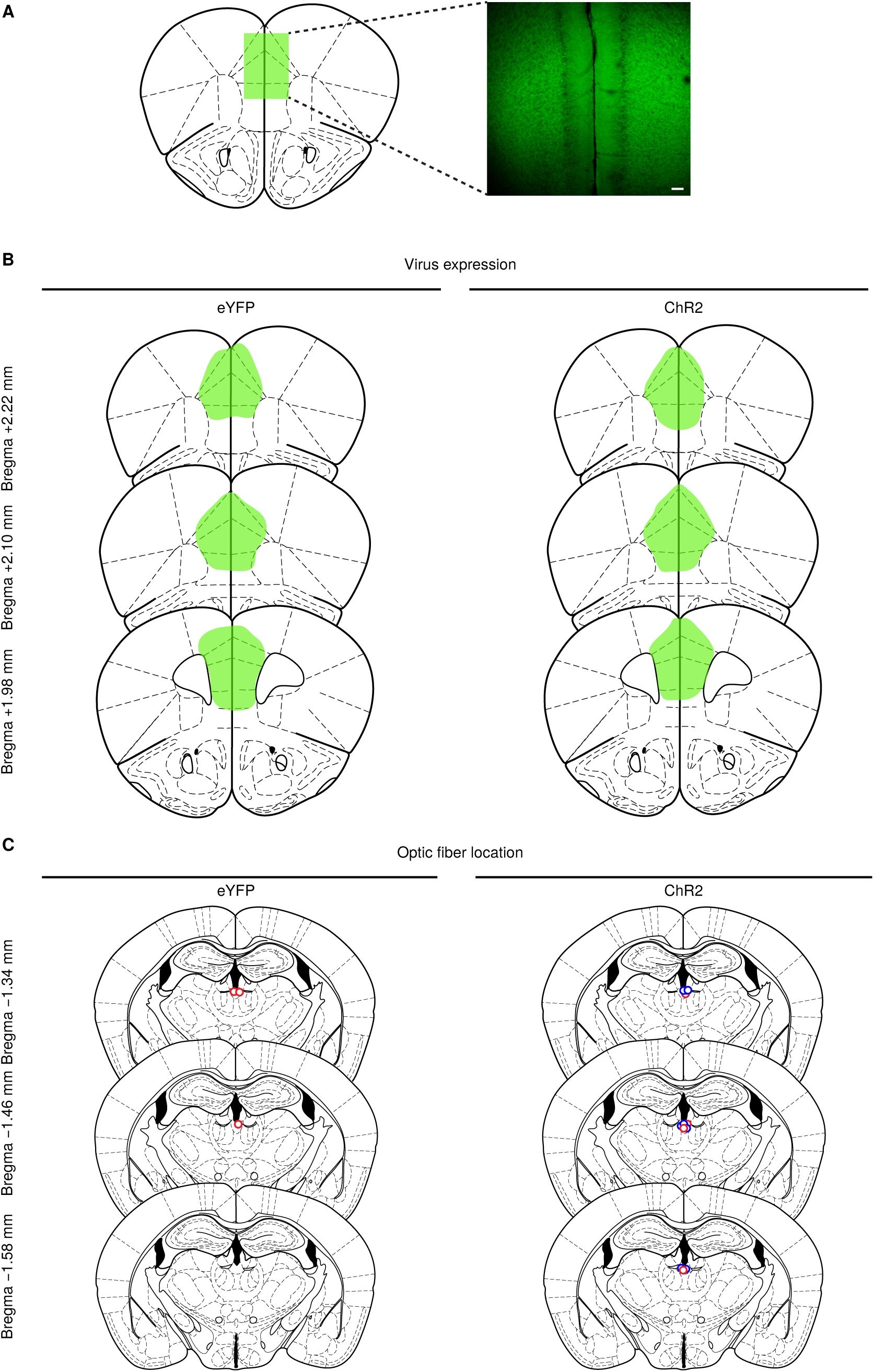
(A) Schematic drawing of viral stereotaxic injection into mPFC. Inset: eYFP (green) expression in a mPFC coronal section from a BL6 mouse that received AAV1-CaMKII*α*-ChR2-eYFP into mPFC (scale bar, 50 *µ*m). (B) Traces showing the expression of eYFP (left) and ChR2-eYFP (right) groups. (C) Locations of fiber tips in eYFP (left) and ChR2-eYFP (right, red=10 Hz and blue=20 HZ) groups in PVT.

**Supplementary Figure 4.**
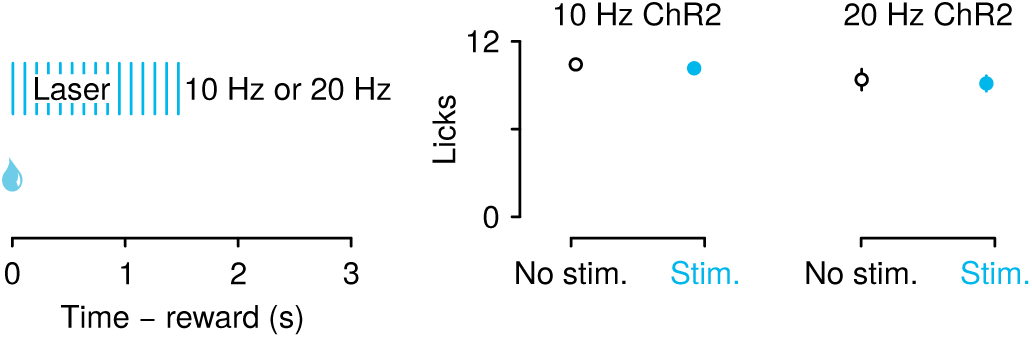
mPFC→PVT stimulation did not suppress licking for unexpected rewards. Left: optical stimulation was delivered during the presentation of reward delivery and lasted for 1500 ms, Right: licking behavior in 10 Hz ChR2eYFP (left) and 20 Hz ChR2-eYFP (right) groups across trials with no stimulation (black) and trials with stimulation (light blue). Line and error bars represent the mean *±* SEM.

## References

Adhikari A, Lerner TN, Finkelstein J, Pak S, Jennings JH, Davidson TJ, Ferenczi E, Gunaydin LA, Mirzabekov JJ, Ye L, Kim SY, Lei A, Deisseroth K. Basomedial amygdala mediates top-down control of anxiety and fear. Nature 527: 179–185, 2015.

Alexander WH, Brown JW. Medial prefrontal cortex as an action-outcome predictor. Nat Neurosci 14: 1338–1344, 2011.

Amemori Ki, Graybiel AM. Localized microstimulation of primate pregenual cingulate cortex induces negative decision-making. Nat Neurosci 15: 776–785, 2012.

Arnsten AFT, Wang MJ, Paspalas CD. Neuromodulation of thought: flexibilities and vulnerabilities in prefrontal cortical network synapses. Neuron 76: 223–239, 2012.

Bach DR, Seymour B, Dolan RJ. Neural activity associated with the passive prediction of ambiguity and risk for aversive events. J Neurosci 29: 1648–1656, 2009.

Bari BA, Grossman CD, Lubin EE, Rajagopalan AE, Cressy JI, Cohen JY. Stable representations of decision variables for flexible behavior. Neuron 103: 922–933.e7, 2019.

Beas BS, Wright BJ, Skirzewski M, Leng Y, Hyun JH, Koita O, Ringelberg N, Kwon HB, Buonanno A, Penzo MA. The locus coeruleus drives disinhibition in the midline thalamus via a dopaminergic mechanism. Nat Neurosci 21: 963–973, 2018.

Bechara A, Damasio H, Damasio A. Emotion, decision making and the orbitofrontal cortex. Cereb Cortex 10: 295–307, 2000.

Beck AT. The evolution of the cognitive model of depression and its neurobiological correlates. Am J Psychiat 165: 969–977, 2008.

Beck CH, Fibiger HC. Conditioned fear-induced changes in behavior and in the expression of the immediate early gene c-fos: with and without diazepam pretreatment. J Neurosci 15: 709–720, 1995.

Bishop P, Burke W, Davis R. Single-unit recording from antidromically activated optic radiation neurones. J Physiol 162: 432–450, 1962.

Blum S, Runyan JD, Dash PK. Inhibition of prefrontal protein synthesis following recall does not disrupt memory for trace fear conditioning. BMC Neurosci 7: 67, 2006.

Boleij H, van’t Klooster J, Lavrijsen M, Kirchhoff S, Arndt SS, Ohl F. A test to identify judgement bias in mice. Behav Brain Res 233: 45–54, 2012.

Browning JR, Jansen HT, Sorg BA. Inactivation of the paraventricular thalamus abolishes the expression of cocaine conditioned place preference in rats. Drug Alcohol Depend 134: 387–390, 2014.

Bubser M, Deutch AY. Stress induces Fos expression in neurons of the thalamic paraventricular nucleus that innervate limbic forebrain sites. Synapse 32: 13–22, 1999.

Burgos-Robles A, Bravo-Rivera H, Quirk GJ. Prelimbic and infralimbic neurons signal distinct aspects of appetitive instrumental behavior. PLoS ONE 8: e57575, 2013.

Burgos-Robles A, Kimchi EY, Izadmehr EM, Porzenheim MJ, Ramos-Guasp WA, Nieh EH, Felix-Ortiz AC, Namburi P, Leppla CA, Presbrey KN, Anandalingam KK, Pagan-Rivera PA, Anahtar M, Beyeler A, Tye KM. Amygdala inputs to prefrontal cortex guide behavior amid conflicting cues of reward and punishment. Nat Neurosci 20: 824–835, 2017.

Burgos-Robles A, Vidal-Gonzalez I, Quirk GJ. Sustained conditioned responses in prelimbic prefrontal neurons are correlated with fear expression and extinction failure. J Neuroci 29: 8474–8482, 2009.

Cerqueira JJ, Mailliet F, Almeida OFX, Jay TM, Sousa N. The prefrontal cortex as a key target of the maladaptive response to stress. J Neurosci 27: 2781–2787, 2007.

Chan SWY, Harmer CJ, Goodwin GM, Norbury R. Risk for depression is associated with neural biases in emotional categorisation. Neuropsychologia 46: 2896–2903, 2008.

Chastrette N, Pfaff DW, Gibbs RB. Effects of daytime and nighttime stress on Fos-like immunoreactivity in the paraventricular nucleus of the hypothalamus, the habenula, and the posterior paraventricular nucleus of the thalamus. Brain Res 563: 339–344, 1991.

Choi DL, Davis JF, Fitzgerald ME, Benoit SC. The role of orexin-A in food motivation, reward-based feeding behavior and food-induced neuronal activation in rats. Neuroscience 167: 11–20, 2010.

Choi EA, Jean-Richard-dit Bressel P, Clifford CW, McNally GP. Paraventricular thalamus controls behavior during motivational conflict. J Neurosci pp. 2480–18, 2019.

Cohen JY, Haesler S, Vong L, Lowell BB, Uchida N. Neuron-type-specific signals for reward and punishment in the ventral tegmental area. Nature 482: 85–88, 2012.

Cook SC, Wellman CL. Chronic stress alters dendritic morphology in rat medial prefrontal cortex. J Neurobiol 60: 236–248, 2004.

Corcoran KA, Quirk GJ. Activity in prelimbic cortex is necessary for the expression of learned, but not innate, fears. J Neurosci 27: 840–844, 2007.

Courtin J, Chaudun F, Rozeske RR, Karalis N, Gonzalez-Campo C, Wurtz H, Abdi A, Baufreton J, Bienvenu TCM, Herry C. Prefrontal parvalbumin interneurons shape neuronal activity to drive fear expression. Nature 505: 92–96, 2014.

Cullinan WE, Herman JP, Battaglia DF, Akil H, Watson SJ. Pattern and time course of immediate early gene expression in rat brain following acute stress. Neuroscience 64: 477–505, 1995.

Darian-Smith I, Phillips G, Ryan R. Functional organization in the trigeminal main sensory and rostral spinal nuclei of the cat. J Physiol 168: 129–146, 1963.

Dearing KF, Gotlib IH. Interpretation of ambiguous information in girls at risk for depression. J Abnorm Child Psychol 37: 79–91, 2009.

Del Arco A, Park J, Wood J, Kim Y, Moghaddam B. Adaptive encoding of outcome prediction by prefrontal cortex ensembles supports behavioral flexibility. J Neurosci 37: 8363–8373, 2017.

Deldin PJ, Levin IP. The effect of mood induction in a risky decision-making task. Bull Psychon Soc 24: 4–6, 1986.

Do-Monte FH, Minier-Toribio A, Quiñones-Laracuente K, Medina-Coln EM, Quirk GJ. Thalamic Regulation of Sucrose Seeking during Unexpected Reward Omission. Neuron 94: 388–400.e4, 2017.

Do-Monte FH, Quiñones-Laracuente K, Quirk GJ. A temporal shift in the circuits mediating retrieval of fear memory. Nature 519: 460–463, 2015.

Dolan R. Emotion, cognition, and behavior. Science 298: 1191–1194, 2002.

Drevets WC, Price JL, Furey ML. Brain structural and functional abnormalities in mood disorders: implications for neurocircuitry models of depression. Brain Struct Funct 213: 93–118, 2008.

Enkel T, Gholizadeh D, von Bohlen Und Halbach O, Sanchis-Segura C, Hurlemann R, Spanagel R, Gass P, Vollmayr B. Ambiguous-cue interpretation is biased under stress- and depression-like states in rats. Neuropsychopharmacology 35: 1008–1015, 2010.

Ernst M, Paulus MP. Neurobiology of decision making: a selective review from a neurocognitive and clinical perspective. Biol Psychiat 58: 597–604, 2005.

Eysenck MW, MacLeod C, Mathews A. Cognitive functioning and anxiety. Psychol Res 49: 189–195, 1987.

Gao C, Leng Y, Ma J, Rooke V, Rodriguez-Gonzalez S, Ramakrishnan C, Deisseroth K, Penzo MA. Two genetically, anatomically and functionally distinct cell types segregate across anteroposterior axis of paraventricular thalamus. Nat Neurosci 23: 217–228, 2020.

Goldwater DS, Pavlides C, Hunter RG, Bloss EB, Hof PR, McEwen BS, Morrison JH. Structural and functional alterations to rat medial prefrontal cortex following chronic restraint stress and recovery. Neuroscience 164: 798–808, 2009.

Haight JL, Flagel SB. A potential role for the paraventricular nucleus of the thalamus in mediating individual variation in Pavlovian conditioned responses. Front Behav Neurosci 8: 79, 2014.

Harding EJ, Paul ES, Mendl M. Animal behaviour: cognitive bias and affective state. Nature 427: 312, 2004.

Haselton MG, Bryant GA, Wilke A, Frederick DA, Galperin A, Frankenhuis WE, Moore T. Adaptive rationality: An evolutionary perspective on cognitive bias. Soc Cogn 27: 733–763, 2009.

Hockey GRJ, John Maule A, Clough PJ, Bdzola L. Effects of negative mood states on risk in everyday decision making. Cogn Emot 14: 823–855, 2000.

Holland PC, Gallagher M. Amygdala-frontal interactions and reward expectancy. Curr Opin Neurobiol 14: 148–155, 2004.

Hsu DT, Kirouac GJ, Zubieta JK, Bhatnagar S. Contributions of the paraventricular thalamic nucleus in the regulation of stress, motivation, and mood. Front Behav Neurosci 8: 73, 2014.

Igelstrom KM, Herbison AE, Hyland BI. Enhanced c-Fos expression in superior colliculus, paraventricular thalamus and septum during learning of cue-reward association. Neuroscience 168: 706–714, 2010.

Ishikawa A, Ambroggi F, Nicola SM, Fields HL. Contributions of the amygdala and medial prefrontal cortex to incentive cue responding. Neuroscience 155: 573–584, 2008.

James MH, Dayas CV. What about me? The PVT: a role for the paraventricular thalamus (PVT) in drug-seeking behavior. Front Behav Neurosci 7: 18, 2013.

Kirouac GJ. Placing the paraventricular nucleus of the thalamus within the brain circuits that control behavior. Neurosci Biobehav Rev 56: 315–329, 2015.

Kukolja J, Schläpfer TE, Keysers C, Klingmüller D, Maier W, Fink GR, Hurlemann R. Modeling a negative response bias in the human amygdala by noradrenergic-glucocorticoid interactions. J Neurosci 28: 12868–12876, 2008.

Lawson C, MacLeod C, Hammond G. Interpretation revealed in the blink of an eye: depressive bias in the resolution of ambiguity. J Abnorm Psychol 111: 321–328, 2002.

Lee JH, Durand R, Gradinaru V, Zhang F, Goshen I, Kim DS, Fenno LE, Ramakrishnan C, Deisseroth K. Global and local fMRI signals driven by neurons defined optogenetically by type and wiring. Nature 465: 788–792, 2010.

Lerner JS, Keltner D. Beyond valence: Toward a model of emotion-specific influences on judgement and choice. Cogn Emot 14: 473–493, 2000.

Levy I, Snell J, Nelson AJ, Rustichini A, Glimcher PW. Neural representation of subjective value under risk and ambiguity. J Neurophysiol 103: 1036–1047, 2010.

Li S, Kirouac GJ. Sources of inputs to the anterior and posterior aspects of the paraventricular nucleus of the thalamus. Brain Struct Funct 217: 257–273, 2012.

Li Y, Lindemann C, Goddard MJ, Hyland BI. Complex Multiplexing of Reward-Cue- and Licking-Movement-Related Activity in Single Midline Thalamus Neurons. J Neurosci 36: 3567–3578, 2016.

Liston C, Miller MM, Goldwater DS, Radley JJ, Rocher AB, Hof PR, Morrison JH, McEwen BS. Stress-induced alterations in prefrontal cortical dendritic morphology predict selective impairments in perceptual attentional set-shifting. J Neurosci 26: 7870–7874, 2006.

Liu RJ, Aghajanian GK. Stress blunts serotonin- and hypocretin-evoked EPSCs in prefrontal cortex: role of corticosterone-mediated apical dendritic atrophy. Proc Natl Acad Sci U S A 105: 359–364, 2008.

Luce RD. Response Times: Their Role in Inferring Elementary Mental Organization. Oxford University Press, 1986.

Luk CH, Wallis JD. Dynamic encoding of responses and outcomes by neurons in medial prefrontal cortex. J Neurosci 29: 7526–7539, 2009.

MacLeod A, Byrne A. Anxiety, depression, and the anticipation of future positive and negative experiences. J Abnorm Psychol 105: 286–289, 1996.

Martin-Fardon R, Boutrel B. Orexin/hypocretin (Orx/Hcrt) transmission and drug-seeking behavior: is the paraventricular nucleus of the thalamus (PVT) part of the drug seeking circuitry? Front Behav Neurosci 6: 75, 2012.

Matzeu A, Weiss F, Martin-Fardon R. Transient inactivation of the posterior paraventricular nucleus of the thalamus blocks cocaine-seeking behavior. Neurosci Lett 608: 34–39, 2015.

Moorman DE, Aston-Jones G. Prefrontal neurons encode context-based response execution and inhibition in reward seeking and extinction. Proc Natl Acad Sci U S A 112: 9472–9477, 2015.

Morgan MA, LeDoux JE. Differential contribution of dorsal and ventral medial prefrontal cortex to the acquisition and extinction of conditioned fear in rats. Behav Neurosci 109: 681–688, 1995.

Opris I, Bruce CJ. Neural circuitry of judgment and decision mechanisms. Brain Res Brain Res Rev 48: 509–526, 2005.

Orsini CA, Heshmati SC, Garman TS, Wall SC, Bizon JL, Setlow B. Contributions of medial prefrontal cortex to decision making involving risk of punishment. Neuropharmacology 139: 205–216, 2018.

Otis JM, Namboodiri VMK, Matan AM, Voets ES, Mohorn EP, Kosyk O, McHenry JA, Robinson JE, Resendez SL, Rossi MA, Stuber GD. Prefrontal cortex output circuits guide reward seeking through divergent cue encoding. Nature 543: 103–107, 2017.

Paintal A. Intramuscular propagation of sensory impulses. J Physiol 148: 240–251, 1959.

Papciak J, Popik P, Fuchs E, Rygula R. Chronic psychosocial stress makes rats more ‘pessimistic’ in the ambiguous-cue interpretation paradigm. Behav Brain Res 256: 305–310, 2013.

Parker RMA, Paul ES, Burman OHP, Browne WJ, Mendl M. Housing conditions affect rat responses to two types of ambiguity in a reward-reward discrimination cognitive bias task. Behav Brain Res 274: 73–83, 2014.

Penzo MA, Robert V, Tucciarone J, De Bundel D, Wang M, Van Aelst L, Darvas M, Parada LF, Palmiter RD, He M, Huang ZJ, Li B. The paraventricular thalamus controls a central amygdala fear circuit. Nature 519: 455–459, 2015.

Peters J, Kalivas PW, Quirk GJ. Extinction circuits for fear and addiction overlap in prefrontal cortex. Learn Mem 16: 279–288, 2009.

Phillips ML, Drevets WC, Rauch SL, Lane R. Neurobiology of emotion perception I: The neural basis of normal emotion perception. Biol Psychiat 54: 504–514, 2003.

Pizzagalli DA, Iosifescu D, Hallett LA, Ratner KG, Fava M. Reduced hedonic capacity in major depressive disorder: evidence from a probabilistic reward task. J Psychiat Res 43: 76–87, 2008.

Radley JJ, Rocher AB, Janssen WGM, Hof PR, McEwen BS, Morrison JH. Reversibility of apical dendritic retraction in the rat medial prefrontal cortex following repeated stress. Exp Neurol 196: 199–203, 2005.

Radley JJ, Rocher AB, Miller M, Janssen WGM, Liston C, Hof PR, McEwen BS, Morrison JH. Repeated stress induces dendritic spine loss in the rat medial prefrontal cortex. Cereb Cortex 16: 313–320, 2006.

Radley JJ, Rocher AB, Rodriguez A, Ehlenberger DB, Dammann M, McEwen BS, Morrison JH, Wearne SL, Hof PR. Repeated stress alters dendritic spine morphology in the rat medial prefrontal cortex. J Comp Neurol 507: 1141–1150, 2008.

Radley JJ, Sisti HM, Hao J, Rocher AB, McCall T, Hof PR, McEwen BS, Morrison JH. Chronic behavioral stress induces apical dendritic reorganization in pyramidal neurons of the medial prefrontal cortex. Neuroscience 125: 1–6, 2004.

Rygula R, Papciak J, Popik P. Trait pessimism predicts vulnerability to stress-induced anhedonia in rats. Neuropsychopharmacology 38: 2188–2196, 2013.

Rygula R, Papciak J, Popik P. The effects of acute pharmacological stimulation of the 5-HT, NA and DA systems on the cognitive judgement bias of rats in the ambiguous-cue interpretation paradigm. Eur Neuropsychopharmacol 24: 1103–1111, 2014.

Rygula R, Pluta H, Popik P. Laughing rats are optimistic. PLoS One 7: e51959, 2012.

Sangha S, Robinson PD, Greba Q, Davies DA, Howland JG. Alterations in reward, fear and safety cue discrimination after inactivation of the rat prelimbic and infralimbic cortices. Neuropsychopharmacology 39: 2405–2413, 2014.

Schiltz CA, Bremer QZ, Landry CF, Kelley AE. Food-associated cues alter forebrain functional connectivity as assessed with immediate early gene and proenkephalin expression. BMC Biol 5: 16, 2007.

Schiltz CA, Kelley AE, Landry CF. Contextual cues associated with nicotine administration increase arc mRNA expression in corticolimbic areas of the rat brain. Eur J Neurosci 21: 1703–1711, 2005.

Schmitzer-Torbert N, Jackson J, Henze D, Harris K, Redish A. Quantitative measures of cluster quality for use in extracellular recordings. Neuroscience 131: 1–11, 2005.

Schwarz N, Clore GL. Mood, misattribution, and judgments of well-being: informative and directive functions of affective states. J Personal Soc Psychol 45: 513, 1983.

Sharp FR, Sagar SM, Hicks K, Lowenstein D, Hisanaga K. c-fos mRNA, Fos, and Fos-related antigen induction by hypertonic saline and stress. J Neurosci 11: 2321–2331, 1991.

Sierra-Mercado D, Padilla-Coreano N, Quirk GJ. Dissociable roles of prelimbic and infralimbic cortices, ventral hippocampus, and basolateral amygdala in the expression and extinction of conditioned fear. Neuropsychopharmacology 36: 529–538, 2011.

Sotres-Bayon F, Quirk GJ. Prefrontal control of fear: more than just extinction. Curr Opin Neurobiol 20: 231–235, 2010.

Sparta DR, Hovelsø N, Mason AO, Kantak PA, Ung RL, Decot HK, Stuber GD. Activation of prefrontal cortical parvalbumin interneurons facilitates extinction of reward-seeking behavior. J Neurosci 34: 3699–3705, 2014.

Spencer SJ, Fox JC, Day TA. Thalamic paraventricular nucleus lesions facilitate central amygdala neuronal responses to acute psychological stress. Brain Res 997: 234–237, 2004.

Stoelzel CR, Bereshpolova Y, Alonso JM, Swadlow HA. Axonal Conduction Delays, Brain State, and Cortico-geniculate Communication. J Neurosci 37: 6342–6358, 2017.

Sugrue LP, Corrado GS, Newsome WT. Choosing the greater of two goods: neural currencies for valuation and decision making. Nat Rev Neurosci 6: 363–375, 2005.

Swadlow HA. Neocortical efferent neurons with very slowly conducting axons: strategies for reliable antidromic identification. J Neurosci Meth 79: 131–141, 1998.

Swadlow HA, Weyand TG. Efferent systems of the rabbit visual cortex: laminar distribution of the cells of origin, axonal conduction velocities, and identification of axonal branches. J Comp Neurol 203: 799–822, 1981.

Vertes RP. Differential projections of the infralimbic and prelimbic cortex in the rat. Synapse 51: 32–58, 2004.

Wei Q, Hebda-Bauer EK, Pletsch A, Luo J, Hoversten MT, Osetek AJ, Evans SJ, Watson SJ, Seasholtz AF, Akil H. Overexpressing the glucocorticoid receptor in forebrain causes an aging-like neuroendocrine phenotype and mild cognitive dysfunction. J Neurosci 27: 8836–8844, 2007.

Wellman CL. Dendritic reorganization in pyramidal neurons in medial prefrontal cortex after chronic corticosterone administration. J Neurobiol 49: 245–253, 2001.

Wright WF, Bower GH. Mood effects on subjective probability assessment. Organiz Behav Hum Decis Proc 52: 276–291, 1992.

Yasoshima Y, Scott TR, Yamamoto T. Differential activation of anterior and midline thalamic nuclei following retrieval of aversively motivated learning tasks. Neuroscience 146: 922–930, 2007.

Yuen EY, Wei J, Liu W, Zhong P, Li X, Yan Z. Repeated stress causes cognitive impairment by suppressing glutamate receptor expression and function in prefrontal cortex. Neuron 73: 962–977, 2012.

Zhu Y, Wienecke CFR, Nachtrab G, Chen X. A thalamic input to the nucleus accumbens mediates opiate dependence. Nature 530: 219–222, 2016.

